# iGABASnFR Imaging Reveals Diffusion-Driven GABA Clearance in the Cerebral Cortex

**DOI:** 10.1101/2025.10.07.680664

**Authors:** Reyna L. Gariepy, Jesse B. Blackman, Jeffrey S. Diamond, Chris G. Dulla, Moritz Armbruster

## Abstract

GABAergic signaling consists of presynaptic release, synaptic and extra-synaptic receptor activation, and signal termination by one of several mechanisms including receptor desensitization, diffusion of GABA, and active clearance by GABA transporters (GATs). Understanding how long GABA is free in the extracellular space is key to understanding how inhibition controls activity but has been technically challenging. Estimates of GABA’s persistence from GABA_A_ and GABA_B_ recordings range from tens of milliseconds to multiple seconds, but may reflect receptor properties rather than extracellular GABA dynamics. Using the fluorescent GABA sensor, iGABASnFr, in the mouse cerebral cortex, we show that GABA rapidly disperses (tens of milliseconds) from sites of release sites via diffusion rather than GATs. This GABA then accumulates in the extracellular space, where GATs require hundreds of milliseconds to remove extracellular GABA following its release, and even longer when the local density of GABA release is elevated. This extracellular summation of GABA can act hetero-synaptically by activating and/or desensitizing both extra-synaptic and neighboring synaptic receptors. Together these findings reveal a stark disconnect between phasic IPSC signals, which rapidly depress, and extracellular GABA, which strongly accumulates, raising new questions about how GABAergic inhibition works to shape network function.

## Introduction

Neurotransmission consists of presynaptic release of neurotransmitter and subsequent receptor activation and is terminated when, depending on the system, neurotransmitter is removed by active transport, diffuses away from the receptor, or is enzymatically degraded. Neurotransmitter receptors also inactivate via desensitization while transmitters are still present resulting in multiple mechanisms that shape how extracellular neurotransmitters dynamically activate receptors. For the excitatory neurotransmitter glutamate, astrocyte excitatory amino acid transporters (EAATs) rapidly bind and clear glutamate from the extracellular space to spatiotemporally limit excitation and keep glutamate levels low^1,2^. The extracellular dynamics of GABA, the primary inhibitory neurotransmitter in the brain, are much less well understood but mediate complex signaling across cell types and at multiple subcellular locations. Low-affinity synaptic GABA_A_ receptors (GABA_A_Rs) mediate phasic inhibition, high-affinity extrasynaptic GABA_A_Rs drive tonic inhibition^3–6^, and metabotropic GABA_B_Rs^4,7^ are coupled to diverse downstream effectors. In addition, local ultrastructure likely shapes extracellular GABA dynamics, so that different GABARs may experience unique local exposure to GABA^8^. GABA transporters (GATs; GAT1, *SLC6A1*, presynaptically expressed by GABAergic neurons; GAT3, *SLC6A11*, expressed by astrocytes)^6,9,10^ actively remove GABA from the extracellular space. GATs are expressed at far lower levels than EAATs^11,12^. GAT1 loss-of-function mutations cause epilepsy and ataxia^13,14^, but pharmacological inhibition of GAT1 with tiagabine is used as an anti-seizure medication^4,14,15^, suggesting complex effects of altering extracellular GABA dynamics. GABAergic inhibition is critical to temporal encoding, information processing, and constraining circuit activity^16,17^ so our lack of understanding of extracellular GABA dynamics represents a significant challenge to decoding inhibitory signaling and related dysfunctions.

The kinetics of phasic inhibition (decay ∼ 20-30 ms) are thought to be largely driven by GABA_A_ receptor properties, primarily desensitization^18–21^. This is evidenced by miniature and spontaneous IPSCs being unaffected by pharmacological inhibition or genetic deletion of GATs. Stimulus-evoked IPSCs, on the other hand, are prolonged by GAT inhibition, suggesting that a single evoked response can lead to GABA diffusion out of the synapse to activate nearby receptors^5,15,22,23^. While this argues for rapid GAT function within the time-scale of an IPSC, measurements of GABA_B_ IPSCs^24,25^, presynaptic effects of GABA_B_^26,27^, GAT transporter current recordings^28^, and modeling^29^ suggest that GABA persists in the extracellular space for 100’s milliseconds to seconds. Consistent with this model, inhibiting GATs enhances tonic GABA inhibition^5,30,31^. Together, this suggests that GABA dynamics are distinct in different extracellular regions, with estimates spanning from milliseconds to seconds based on limited tools to measure quantitatively extracellular GABA in space and time.

Previously, we have employed iGluSnFR, the fluorescence glutamate sensor, to quantify extracellular glutamate clearance^32,33^. Here we use iGABASnFR, a fluorescent intensity-based GABA sensor^34,35^, to quantify extracellular GABA dynamics in the mouse cortex. Using epifluorescence imaging, which reports spatially averaged GABA in the extracellular space, we report that following synaptic stimulation, GABA persists in the extracellular space for 100’s of milliseconds with GATs modestly contributing to GABA clearance. Using modeling and high resolution confocal imaging to investigate GABA surrounding release sites, we show that GABA disperses rapidly into the extracellular space with minimal contribution from GAT activity. This local elevation of GABA, even for small bursts of synaptic activity, can spillover onto neighboring synapses and drive heterosynaptic suppression of inhibition, presumably via receptor desensitization. This suggests that stimulus-evoked GABA release is largely a volumetric signal with GABA diffusing from the synapse even for a single stimulus and activating both synaptic and extrasynaptic receptors, blurring the distinction between phasic and tonic GABA.

## Methods

### Animals

All procedures were approved by the Tufts University Institutional Animal Care and Use Committee. Male and female C57/Bl6 mice were bred in house from mice originally obtained from Jackson Labs (Strain: 000664), kept on a 12-h light/dark cycle with access to food and water ad libitum.

### Adeno-associated virus injection

Somatosensory cortical expression of the iGABASnFr1 or iGABASnFr2 was attained via focal viral injection of iGABASnFr1^34^ (AAV1.hSyn.iGABASnFR, Addgene: 112159) or iGABASnFr2^35^ (AAV.php.eB.hSyn.iGABASnFR2-WPRE, Virovek). For iGluSnFr imaging AAV5-hSyn-iGluSnFr^36^ (University of Pennsylvania Vector Core; catalog #AV-5-PV2723, AV-5-PV2914) was injected as previously described^32^. C57/BL6 mice were stereotactically injected with 3 injection sites in a single hemisphere (coordinates): (1.25, 1.25, 0.5), (1.25, 2.25, 0.5), and (1.25, 3.25, 0.5) (λ + x, +y, −z) mm, 1µl volume was injected per site with ∼5E9 vector genomes. Following injection, mice were monitored closely for 3 days to ensure recovery. Acute brain slices were prepared for imaging 2-4 weeks following injection. We would like to thank Dr. Loren Looger and the GENIE Project for making the plasmids and viruses available that were used in this study.

### Preparations of acute brain slices

Acute brain slices were prepared as described previously^32^. Mice were anesthetized with isoflurane, decapitated, and the brains were rapidly removed and placed in ice-cold slicing solution containing (in mM): 2.5 KCl, 1.25 NaH_2_PO_4_, 10 MgSO_4_, 0.5 CaCl_2_, 11 glucose, 234 sucrose, and 26 NaHCO_3_ and equilibrated with 95% O_2_:5% CO_2_. The brain was glued to a Vibratome VT1200S (Leica Microsystems, Wetzlar, Germany), and slices (400 μm thick) were cut in a coronal orientation. Slices were then placed into a recovery chamber containing aCSF comprising (in mM): 126 NaCl, 2.5 KCl, 1.25 NaH_2_PO_4_, 1 MgSO_4_, 2 CaCl_2_, 10 glucose, and 26 NaHCO_3_ (equilibrated with 95% O_2_:5% CO_2_). Slices were allowed to equilibrate in aCSF at 32°C for 1 hr.

### Live slice imaging

Slices were placed in a submersion chamber (Siskiyou), held in place with small gold wires, and perfused with aCSF containing 20µM DNQX (antagonist of AMPA receptors, Sigma), and 50µM APV (antagonist of NMDA receptors, Tocris), equilibrated with 95% O_2_: 5% CO_2_ and circulated at 2ml/min at 34°C. A tungsten concentric bipolar stimulating electrode (FHC; Bowdoin, ME USA) was placed in the deep cortical layers and stimulus intensity was set for 2x the minimally detectable stimulus. iGABASnFr1 responses were imaged with a 10X water-immersion objective (Olympus) on a custom Olympus microscope with 470nm LED illumination (Thorlabs). Responses were captured with a Andor Zyla 5.5 (Andor) camera imaging, 12bit digitization, 10ms rolling shutter mode for 100Hz temporal resolution controlled by MicroManager^37^ in epifluorescence mode using a GFP filter cube (Chroma). Stimulus responses were imaged in sequence of 1 stimuli, 5 stimuli at 100Hz, 10 stimuli at 100Hz, and no stimulus bleaching control with 10s between acquisitions. This sequence was repeated 10 times for each slice. For each stimulus, a 3s imaging window was acquired. Drugs to inhibit GAT1 (25 μM NO-711, Abcam: ac120364) and GAT3 (50 μM SNAP-5114, Sigma S1069) were applied for 5 minutes before recommencing imaging.

For confocal imaging and deconvolution we utilized the iGABASnFr2 sensor which has increased affinity and improved sensor performance^35,38^. For iGABASnFr2 confocal imaging, slices were placed in a submersion chamber (Warner Instruments) of a custom Prior Open-scope microscope equipped with an X-light V2 Spinning disk confocal (Crest-optic, 89North LDI). Responses were imaged onto a Prime95B (Photometrics) camera with the same stimulus protocol as described for epifluorescence imaging. Responses were imaged with a 60x water immersion objective (Olympus) with a 2x magnifier at a 100Hz frame rate.

### IPSC recordings

IPSCs were recorded similar to previous studies^39^. Briefly, whole-cell patch-clamp recordings were made in an identical setup to the live-slice imaging, with aCSF containing DNQX and APV. Pyramidal neurons were identified by morphology in layer V of the cortex, whole-cell patch-clamp recordings using 2-5MΩ borosilicate glass electrodes were established using an internal solution containing the following: CsMs 140, HEPES 10, NaCl 5, EGTA 0.2, QX314 5, MgATP 1.8, NaGTP 0.3 in mM, pH 7.25, Osmolarity 290. In voltage-clamp mode, neurons were held at 0 mV (reversal potential for glutamate receptors) to isolate IPSPs, and then IPSCs were evoked by electrical stimulation identically to the live-cell imaging above.

### Deconvolution

Averaged, normalized 10 stimuli 100Hz iGABASnFr1 and iGABASnFR2 traces were used to deconvolve out the sensor properties from the underlying GABA kinetics. We assume that the recorded fluorescence signal is a convolution of the biological time course of GABA in the extracellular space and the iGABASnFr response function, being a function of the Kon and Koff rates:

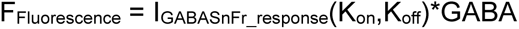

We assume the underlying GABA kinetic is the same independent of which sensor is expressed. Based on published data we assume the affinity of the sensors are iGABASnFr1 K_d_ = 30µM^34^ and iGABASnFr2 K_d_ = 6.4µM^35^. Using the relation that Kd is the balance of *K_off_/K_on_* and we can reduce the variables to 3, *K_on_iGABASnFr1_*, *K_on_iGABASnFr2_*, and the Response function amplitude *A*. Using the Matlab (Mathworks), *fmincon* optimization function to minimize the difference between the iGABASnFR1 GABA response and iGABASnFR2 GABA response, resulting in the iGABASnFr response functions (Fig. S2B). The response functions were then validated on a second dataset which showed matching deconvolved GABA responses from iGABASnFr1 and iGABASnFr2 recordings and applied to 1, 5 and 10 stimuli at 100Hz traces(Fig. S2C).

### Modeling

The GABA diffusion model was adapted based on previous modeling of glutamate diffusion^33^ written in MATLAB. Briefly, the model simulates a single synapse, and a single stimulus GABA release event, releasing 5000 independent GABA molecules that originate from a point source in a 320nm diameter synaptic cleft. For each 1µs time step, each GABA molecule diffuses randomly, with a diffusion coefficient of 0.253µs^2^/ms. For the synaptic cleft, diffusion is restricted to two dimensions, but unrestricted once the molecule leaves the synaptic cleft. Outside the synaptic cleft, diffusion was scaled by the extracellular volume fraction (0.21) and the extracellular tortuosity (λ=1.55). Compared to the previous glutamate diffusion models, GABA transporter density (∼1µM) was reduced 100-fold compared to glutamate transporters (100µM). This is based upon estimates of 800-1300 per µm^2^ GAT-1 molecules at boutons with additional reductions due to intracellular localization of 40% of GATs and reduced expression in other cell types^11,40^. For simplicity we assume homogenous distribution of transporters in the extracellular space. GAT affinity and transport kinetics are very similar to GLT-1 (4-20uM GAT1^41,42^, 12uM GLT-1^43,44^ and 70-160ms GAT1 ^42^vs. 20-70ms GLT-1^43,45^, respectively) so were left unchanged from previous models.

### Analysis

Data was analyzed using MATLAB (Mathworks) and Origin (Originlab). For low magnification (10X) imaging data, the central quadrant (25%-75%) of the camera field of view was isolated, spatially averaged down-sampled 10-fold to reduce data size, and averaged across sweeps. ΔF/F traces were created using the 1, 5, and 10 stimuli traces divided by the no stimulus trace. Due to different bleaching kinetics, the first sweep was excluded. For epifluorescence traces an additional mono-exponential bleaching correction was applied (as needed) based on the pre-stimulus frames and frames at the end of the 3s acquisition.

iGABASnFr decay times were measured using the T_80_-T_20_ time from the end of the stimulus. Traces were normalized 0-1, from the end of the stimulus. Traces were linearly extrapolated 10-fold, and 80% and 20% times were found. For analysis of peak versus decay, for the average ΔF/F image, a signal to noise image was created (imaging stack divided by the prestimulus standard deviation). Peak signal to noise across the imaging window was binned with a bin size of 3. For each bin an average trace was created with peak and decay quantified.

For confocal data, the individual sweeps were aligned with NoRMCorre algorithm^46^. Following averaging, stimulation responses were divided by the no stimulus traces for bleaching correction after subtracting the dark noise and auto fluorescent counts to create ΔF/F_0_ datasets. To establish ROIs from confocal iGABASnFr2 imaging, aligned fluorescence images were averaged and smoothed with a 1.5-pixel 2D gaussian and a one-sided gaussian in the temporal dimension (sigma = 50ms). Next the images were divided by a 15-pixel radius spatial filtered version of the data (pixel size = 92nm) to highlight responsive areas over the background. The two frames post-stimulation were averaged and all responses >4 standard deviations over the baseline, >6 pixels and a circularity of >0.75 were defined as responsive ROIs. ROI masks were then applied to the 1, 5, and 10 stimuli at 100Hz ΔF/F traces.

### Statistics and reproducibility

Individual statistical tests are noted throughout the manuscript and figure legends. Linear mixed models were used whenever possible to account for potential non-independence of slices coming from the same mouse. We used the *lmer* function in R to fit linear mixed models. For all models, statistical significance was defined as alpha < 0.05. When significant main effects or interactions were found, paired or two-sample t-tests or two-way repeated-measures ANOVA with Tukey’s post-test were made to determine differences between specific groups as needed. Sample sizes (slices/cells) are listed for each experiment in the figure labels or text, all experiments are from ≥3 mice, with exact n values for mice, slices, and imaging fields of view noted throughout in the figure legends. Error bars indicate standard error of the mean (SEM).

## Results

### Measuring extracellular GABA dynamics in the cerebral cortex using iGABASnFr imaging

Many studies have attempted to use electrophysiological recording of GABA receptor (GABAR) activity^13,30,47^, transporter currents^28^, and other indirect approaches to infer extracellular GABA dynamics^23–26,29^, each of which come with technical caveats. iGABASnFr imaging enables direct visualization of GABA dynamics in the extracellular space, providing a rapid, optical readout of extracellular GABA. We expressed the neuronally targeted GABA sensor, iGABASnFR1, in layer IV/V of the somatosensory cortex in 4–6-week-old mice via intracortical AAV injection (AAV1-hsyn-iGABASnFR1) (Fig. 1A). Two to four weeks following infection, acute brain slices were prepared for iGABASnFR imaging. GABA release was induced via local electrical stimulation (2X threshold stimulation) using a bipolar stimulating electrode placed in layer V and iGABASnFr1 ΔF/F responses were imaged (10X magnification) in the presence of AMPA and NMDA receptor antagonists (DNQX 20μM and APV 50μM). Spatially averaged fluorescence responses over showed a stimulation-dependent increase in iGABASnFr1 fluorescence immediately upon stimulation (Fig. 1B). iGABASnFR ΔF/F peak significantly increased with increased numbers of stimuli (1, 5, and 10 stimuli delivered at 100Hz; 0.15 ± 1.2E^−4^, 0.44 ± 3.2E^−4^, 0.65 ± 4.1E^−4^ ΔF/F (%), respectively; N= 21 slices, 7 mice) showing the ability to quantify extracellular GABA across a broad dynamic range (Fig. 1C).

**Figure 1:**
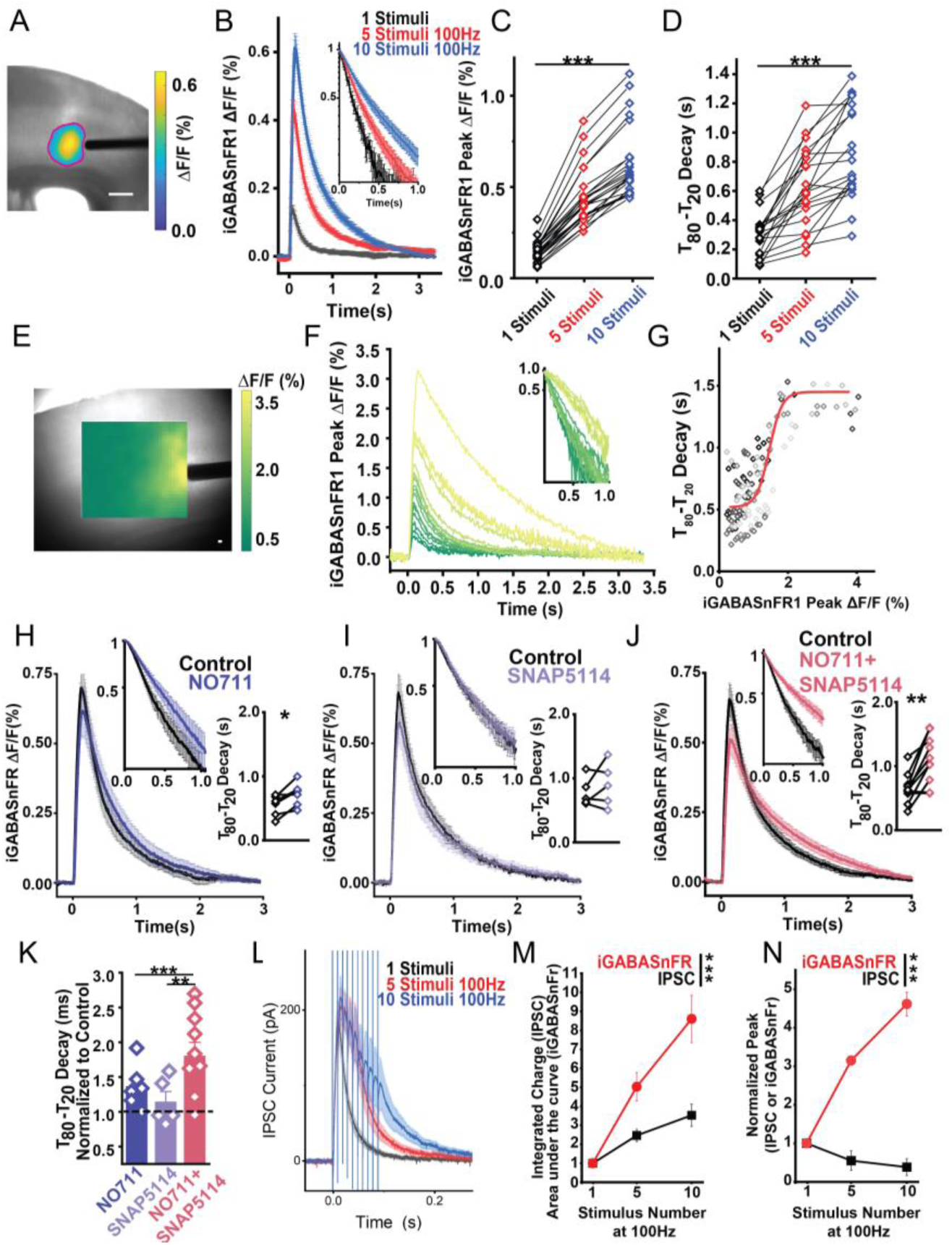
iGABASnFr decay is slow and modulated by local GABA load. **A)** 4x brightfield image with pseudcolored ΔF/F iGABASnFR1 response to 10 stimuli at 100Hz. Scale bar 125μm. **B)** Average iGABASnFR1 ΔF/F traces in response to 1, 5, and 10 stimuli at 100Hz with a semi log inset of the normalized fluorescent decay trace. **C)** iGABASnFR1 peak ΔF/F (%) increases with stimuli number. LMM, stimuli dependence p < 2e^−16^. **D)** T_80_-T_20_ Decay of iGABASnFr1 traces shows stimulus dependence LMM, stimuli dependence p < 7.9e^−13^. N= 7 mice; 21 slices. **E)** Brightfield image with iGABASnFR1 responsed binned by their signal to noise and pseudocolored for 10 stimuli at 100Hz. **F)** Representative example of a single slice’s DF/F response for each bin on the matching color scale. **G)** Average binned iGABASnFr1 peak and decay in response to 10 stimuli at 100Hz for 10 slices fitted with a sigmoidal curve, individual slices color coded. N= 6 mice; 9 slices. **H)** Average traces of iGABASnFR1 in response to 10 stimuli at 100Hz while inhibiting GAT1 with in 25μM NO711, **I)** GAT3 with 50μM SNAP5114, and **J)** both GAT1 and GAT3, with a semi log inset of the normalized fluorescent decay trace and a line series inset of the T_80_-T_20_ Decay (s). Paired t-test (T_80_-T_20_ Decay) NO711, p<0.017 N= 3 mice, 6 slices, SNAP5114 p= 3 mice, 5 slices, NO711+SNAP5114 p<0.002 N= 10 slices, 6 mice. **K)** iGABASnFr decay times for each slice normalized to control shows the additive effect of GAT1+GAT3 inhibition, One way ANOVA Tukey Test NO711 vs NO711+SNAP5114 p<6.8e^−4^, SNAP5114 vs NO711+SNAP5114 p=0.002. **L)** Stimulus evoked IPSCs recorded from Layer V pyramidal neurons, for 1, 5, and 10 stimuli at 100Hz shows much faster decays and reduced summation across stimuli. **M)** IPSC integrated charge plotted against iGABASnFr1 area under the curve shows significantly greater summation of stimuli for iGABASnFr1 compared to IPSCs (LMM p < 2.7E-4). **N)** IPSC peaks show depression with stimuli number compared to summation for iGABASnFr1. LMM p < 2E-16 N=13 cells from 4 mice, N=21 slices from 7 mice. * = p<0.05, ** = p<0.01, *** = p<0.001

### Local GABA levels are correlated with the duration of extracellular GABA persistence

iGABASnFr imaging enabled us to quantify the lifetime of extracellular GABA and how this is shaped by the number of stimuli and the local GABA concentration. iGABASnFr1 fluorescent traces had prolonged decay times (quantified by the T_80_-T_20_ decay time) with increasing numbers of stimuli (Fig. 1B inset, 1D; 299 ± 33ms, 628 ± 61ms, 854 ± 72ms for 1, 5, and 10 stimuli at 100Hz, respectively; N=20 slices, 7 mice). To determine whether the slowed decay times were associated with increased GABA levels, we correlated local iGABASnFR1 peak ΔF/F with T_80_-T_-20_ decay times for each stimulus. iGABASnFr1 ΔF/F responses to 10 stimuli delivered at 100Hz were spatially binned based on its signal to noise. For each bin, the peak ΔF/F and decay times were quantified, enabling us to ask how the local peak response is related to the decay at the same location (Fig. 1E-G). The iGABASnFR peak ΔF/F showed a sigmoidal relationship with decay time (Minimum: 526 ± 33ms, Maximum:1354 ± 54ms, Half max ΔF/F (%): 1.5 ± 0.063, slope: 0.21 ± 0.057, R^2^ = 0.669, Fig. 1G), suggesting that the local extracellular GABA level influences how long GABA persists in the extracellular space and saturates GATs at areas of high focal GABAergic activity. The upper end of the decay time is similar to observed time-constants for dye-diffusion in the cortex^48^, potentially suggesting overwhelmed transport capacity and diffusion-driven decay.

### GATs play a small but significant role in removing extracellular GABA in the cortex

GAT1 and GAT3 are both expressed in the somatosensory cortex^10,49^, with GAT1 primarily expressed in presynaptic neurons and GAT3 primarily expressed in astrocytes^6,50–52^. Loss of either transporter induces changes in network activity and pathologies including epilepsy and ataxia^53,54^. To determine the contribution of each transporter to GABA’s persistence in the extracellular space, we sequentially inhibited GAT1 (25µM NO-711^55^), GAT3 (50µM SNAP-5114^31,56^), and both while quantifying iGABASnFR decays (Fig. 1H-K). Inhibiting GAT1 with NO-711 significantly slowed iGABASnFR1 T_80_-T_20_ decay time (0.55 ± 0.07 vs 0.72 ± 0.08 s, N= 6 slices. Paired sample t-test, p=0.0165, Fig. 1H). Inhibiting GAT3 with SNAP-5114 showed no significant effect on the decay time (0.78 ± 0.1 vs 0.9 ± 0.15 s N= 5 slices. Paired sample t-test, p= 0.39 Fig. 1I). Inhibiting both GAT1 and GAT3 showed a significantly bigger effect, normalized to control, than inhibiting GAT1 alone (Fig. 1J/K; Ratio of decay to control: NO711 1.35 ± 0.13, SNAP5114 1.14 ± 0.15, NO711+SNAP5114 1.80 ± 0.20, One-Way ANOVA Tukey Test, NO711 vs NO711+SNAP5114 p<6.8e-4, SNAP5114 vs NO711+SNAP5114 p=0.002, NO711 vs SNAP514 p=0.63, N=10 slices). Similar results were seen for 1 and 5 stimuli (Fig. S1). This suggests a smaller role for GAT3 compared to GAT1. GAT inhibition also decreased iGABASnFR1 peak ΔF/F (Fig. S2). By comparison, when glutamate transporters are inhibited with the EAAT inhibitor 1µM TFB-TBOA, iGluSnFr shows a large increase in fluorescence (Fig. S3) due to fast glutamate uptake during the stimulus train. To test if changes in peak ΔF/F are due to photobleaching, we plotted the average iGABASnFR peak for each sweep, for each condition (control, NO-711, NO711+SNAP5114) (Fig. S2B). We observed a linear decrease in peak ΔF/F fluorescence per sweep that is not affected by GAT inhibition. This suggests the changes are largely due to photobleaching rather than changes in release. Together this shows that GATs play a small but significant role in removing GABA from the extracellular space.

### Deconvolution of iGABASnFR

iGABASnFr1 imaging shows a very slow decay in response to brief bursts of stimuli, but this could be caused by iGABASnFR kinetics and buffering properties, both of which can artificially prolong the measured kinetics of the underlying GABA signal^1,33,57^. To address these potential confounds, we first used a mathematical deconvolution approach to isolate the underlying GABA transients. We collected GABA imaging data using both iGABASnFR1 and the higher affinity fluorescent GABA sensor, iGABASnFr2^35^ (30 μM, 6 μM affinity for iGABASnFr1 and iGABASnFr2 respectively). By using two different affinity sensors (Fig. S4) and making the assumption that their kinetics (K_on_/K_off_) scale with their affinities^57^, we were able to deconvolve out the filtering effect of the sensors to estimate the underlying GABA kinetics (Fig. S4B). For 10 stimuli delivered at 100 Hz, the deconvolved GABA signals had similar decays to the raw iGABASnFr (Fits on average traces: T_80-20_ iGABASnFr1: 1104 ms vs iGABASnFr1_deconvolved 1196 ms and iGABASnFr2: 978ms vs iGABASnFr2_deconvolved 1084ms), suggesting a minimal effect of filtering and supporting the long persistence of GABA in the extracellular space. Interestingly, for 1 and 5 stimuli the deconvolved GABA signal shows a small rapidly decaying component (Fig. S1C), suggesting GATs can provide some fast GABA clearance. To test if differences in iGABASnFr expression level artificially prolong extracellular GABA dynamics via buffering, we asked if decay time is correlated with the baseline iGABASnFR fluorescence prior to stimulation, a proxy for sensor expression (Fig. S5). For both individual and pooled slices, the baseline fluorescence showed no correlation with the decay (R-squared = 0.002)(Fig. S5). This suggests that, within our experimental regime, the decay kinetics are not substantially altered by iGABASnFr1 expression. Both controls validate our iGABASnFr imaging approach and suggest that the iGABASnFr1 kinetics are a suitable proxy for quantifying GABA dynamics in the extracellular space.

### IPSC integration diverges from extracellular GABA levels

With GABA persisting in the extracellular space for an extended time following release and summating with each stimulus (Fig 1B), we asked how neuronal GABA IPSCs summate for similar stimuli. To compare iGABASnFR1 responses to synaptic inhibition, we performed whole-cell patch clamp recordings from layer V pyramidal neurons and isolated inhibitory post-synaptic currents (IPSCs) by holding the cell at the glutamate receptor reversal potential (0 mV) while blocking AMPA and NMDA receptors. As with iGABASnFR1 imaging, GABA release was induced via local stimulation with a bipolar electrode (Fig. 1L). iGABASnFr1 imaging showed significantly greater summation from 1 to 5, and 10 stimuli compared to IPSCs (iGABASnFr1 Area under the curve: 5.0 ± 0.7 and 8.6 ± 1.3 fold increase of over 1 stimuli for 5 and 10 stimuli respectively N=21 slices from 7 mice, IPSC integrated charge: 2.5 ± 0.32 and 3.5 ± 0.6 fold increase over 1 stimuli for 5 and 10 stimuli. N= 13 cells from 4 mice; LMM p=0.0002) (Fig. 1M). Similarly, IPSC peak at the end of the stimulus train depresses with stimuli number, while iGABASnFr peaks summate (iGABASnFr1 peaks (normalized to 1 stimuli), 3.2 ± 0.1, 4.6 ± 0.3 for 5 and 10 stimuli. IPSC peaks (normalized to 1 stimuli) 0.55 ± 0.25, 0.38 ± 0.22 for 5 and 10 stimuli LMM p < 2E-16, iGABASnFr N=21 slices from 7 mice, IPSC N=13 cells from 4 mice; Fig. 1N). This suggests that each burst of stimulus-induced GABA release can activate fewer GABA_A_ receptors, possibly due to receptor desensitization, even with increasing amounts of GABA in the extracellular space.

### Modeling synaptic GABA clearance

Epifluorescence iGABASnFR1 imaging showed very slow extracellular GABA clearance. This is a strong contrast to glutamate clearance, which is removed from the extracellular space by EAATs in just a few milliseconds ^1,32^ and is seemingly contrary to the effect of GAT1 inhibition on evoked IPSCs^5,15,22,23^. To understand the role of GATs uptake versus diffusion and its effects on synaptic GABA dynamics, we first utilized a computational model. We adapted a Monte Carlo simulation of synaptic glutamate release, diffusion, and clearance^33^ to model iGABASnFr, GATs, and GABA diffusion in the extracellular space. We reduced the concentration of transporters in the model from 100 μM (glutamate)^12^ to 1 μM (GABA) based on experimental estimates^11^. We defined 3 regions of interest (ROIs) in the model, the synaptic space (within 1 μm of the release site), the peri-synaptic (1-3 μm of the release site) and the full field (1000 μm^3^) (Fig. 2A). Simulated iGABASnFR2 (to match subsequent confocal imaging) responses in the synaptic ROI showed a rapid rise, increased peak, and faster decay compared to the full field, with the peri-synaptic ROI showing intermediate properties (Fig. 2B-C). To test the role of GAT-mediated uptake in our simulation, we reduced GAT activity between 30% and 90% by reducing the number of transporter present. This significantly slowed the simulated iGABASnFR2 clearance for the full field but had no effect on synaptic GABA clearance (Fig. 2D-E), suggesting that diffusion rather than transport dominates in the synaptic space. Plotting the cumulative transport of GATs in this model shows that GATs bind a small amount of GABA within the time-course of an IPSC (<30ms), consistent with the effects on IPSCs (Fig. S6), although this accounts for <10% of the total GABA clearance. Together, this suggests that GABA diffuses rapidly out of the synaptic ROI through the perisynaptic space and is then slowly cleared by the GABA transporters.

**Figure 2:**
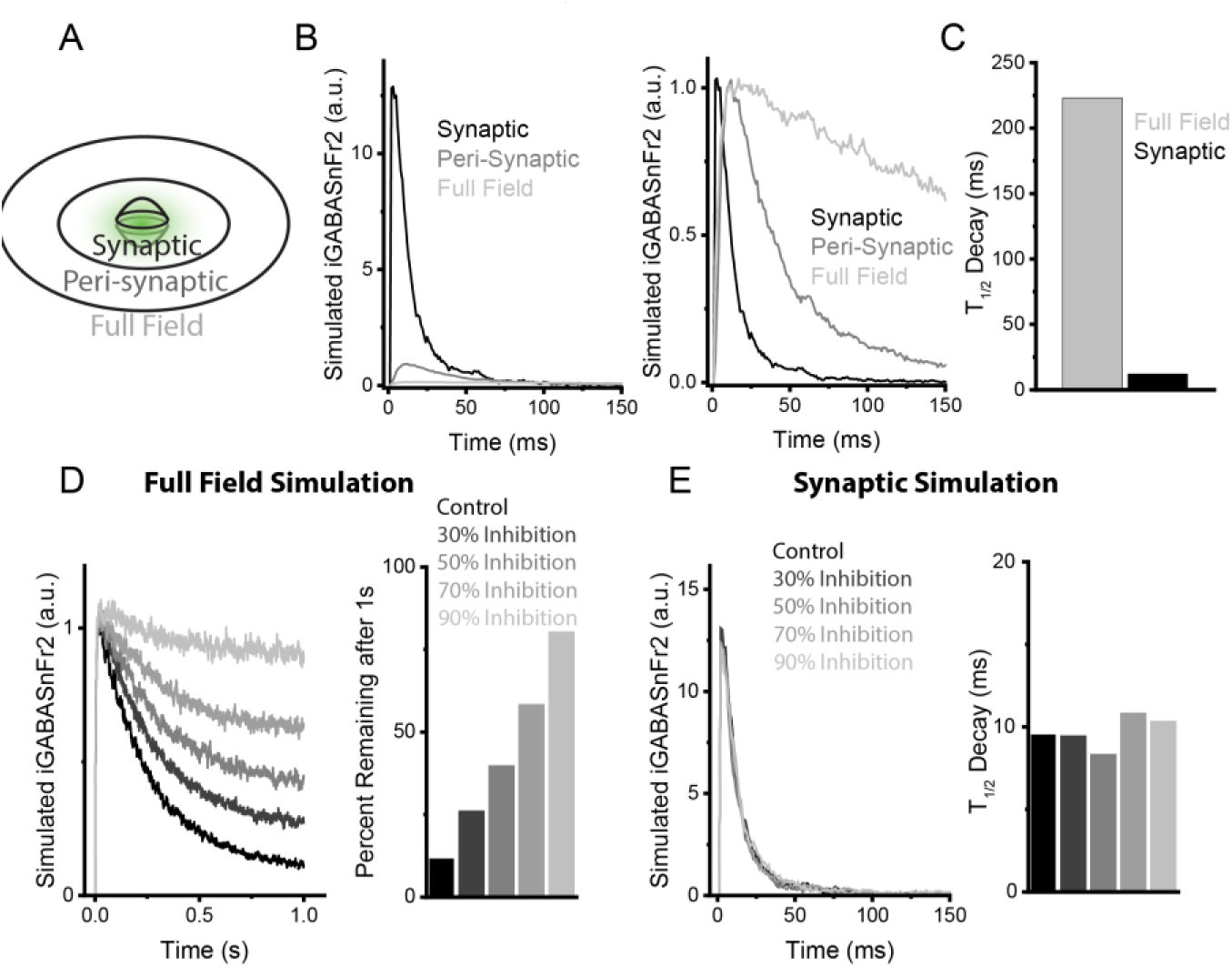
Monte-Carlo modeling of synaptic GABA clearance. **A)** Visualization of a Monte-Carlo diffusion/clearance model of GABA in the extracellular space. GABA is released in a 320nm synaptic cleft, and we have defined 3 ROIs (synaptic <1um, peri-synaptic 1-3um and full field). **B)** Simulated iGABASnFr2 traces of synaptic, peri-synaptic, and full field ROIs both raw and normalized plots. The traces show faster iGABASnFr2 decays in the synaptic ROI compared to the full field, quantified in **C)**. **D)** Model simulations with varying percent inhibition of GATs (30-70%) to simulate GAT inhibitors, shows slowed clearance for full field ROIs, with greater percent of simulated iGABASnFr2 signal remaining 1s after stimulus. **E)** Synaptic ROI shows no effect of GAT inhibition on the clearance kinetic.

### Confocal imaging reveals rapid GABA diffusion from release sites to the extrasynaptic space

To determine if the computational model’s predictions were accurate, we used spinning disk confocal iGluSnFR2^35^ imaging, a higher-affinity and improved sensor, to visualize sites of GABA release. iGABASnFR2 fluorescent responses were repeatedly imaged for a single stimulus and sites of release were identified as small areas that rapidly increased ΔF/F (within 10-20 ms) upon stimulation (Fig. 3A). Similar to modeling results, iGABASnFR2 imaging showed a significantly larger peak around the site of release (Synaptic ROI), as compared to the full field (Fig. 3B) (Peak ΔF/F 4.8E-4, 2.2E-02 for Full Field vs. Synaptic ROI, LMM, p=8.5E-16, from 18 imaging fields from 11 slices from 3 mice). Additionally, the synaptic ROI showed iGABASnFR ΔF/F had a significantly faster decay time as compared to the full field (Fig. 3C; 259 ± 43 ms and 90 ± 12 ms for Full Field vs Synaptic, LMM, p=0.0015 from 18 imaging fields from 11 slices from 3 mice), again mirroring the computational findings. Inhibiting GAT-1 and GAT-3 with NO-711 and SNAP-5114 significantly increased iGABASnFR2 decay times for the full field ROI, but had no significant effect on the decay time of iGABASnFR signal in the Synaptic ROI (Fig. 3D/E) (LMM, Full Field ROI: Control, 242±20 ms, NO-711/SNAP-5114, 400±23 ms, p=5.3E-7; Synaptic ROI: Control, 79±6 ms, NO-711/SNAP-5114 68±8 ms, p=0.29). Modeling suggested a small amount of GAT-mediated GABA clearance occurs during the timescale of an IPSC. Consistent with this, there was a small, but significant, role for GAT clearance in the synaptic ROI seen as an increase in iGABASnFR2 ΔF/F peak when GATs were inhibited (Fig. 3F, LMM, peak ΔF/F (%), Control: 0.11±0.04, NO-711/SNAP-5114: 0.13±0.04, p=0.0015). This increase in peak reflects GAT activity within the duration of an IPSC (Fig. 3F). For single stimuli, the Synaptic ROI had a significantly larger peak than in the full field. If GABA diffuses out of the Synaptic ROI, we would predict that this enrichment would be reduced for larger number stimuli as GABA accumulates in the extracellular space. As predicted, the enrichment of synaptic iGABASnFr2, compared to full field ΔF/F was significantly reduced with stimuli number (Fig. 3G/H; LMM, Stimulus p < 1.4 E-07, from n =16 imaging fields (control and inhibitor) from 11 slices from 3 mice). These results agree with computational findings and suggest that GABA rapidly diffuses out from the synapse where it accumulates in the extrasynaptic space during repeated stimulation, with only a small role for GAT-mediated clearance.

**Figure 3:**
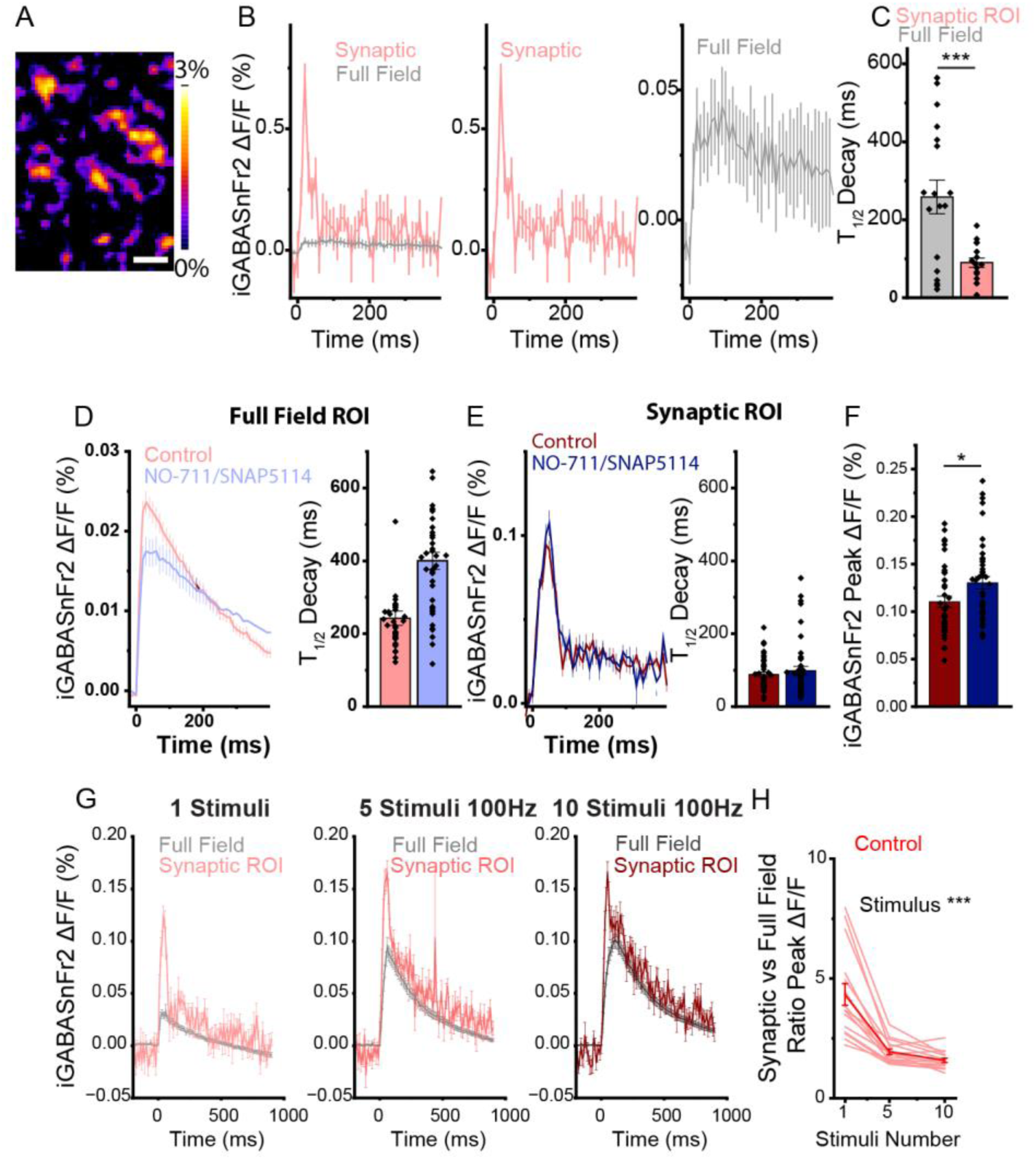
Confocal imaging of synaptic GABA clearance reveals role of diffusion. **A)** Spinning disk confocal imaging of iGABASnFr2 in response to 1 stimulus with release site ROIs highlighted. Scale bar = 1 μm. **B)** Average synaptic, peri-synaptic, and full field confocal iGABASnFr2 traces show faster iGABASnFr2 decays quantified in **C**. LMM p=0.0015, N= 18 imaging fields from 11 slices from 4 mice. **C)** Confocal imaging of iGABASnFr2 with or without NO711 and SNAP5114 shows slowed decay for the full field ROI p=5E-07. **D)** The same imaging for Synaptic ROIs shows no effect on the decay with NO711/SNAP5114 inhibition. p=0.29. N=43, 44 imaging field, from 14 slices from 4 mice. **E)** Synaptic ROIs reveal a small change in peak iGABASnFr2 signal, representing fast GABA clearance at synapses. LMM p=0.0263 N=43,44 imaging fields from 14 slices, 3 mice. **F)** Synaptic compared to Full Field average iGABASnFr2 traces, comparing the enrichment of iGABASnFr2 signal at synapses. **G)** Increasing stimuli number reduce the difference between synaptic and full field iGABASnFr2 traces **H)** Ratio of peak fluorescence (Synaptic ROI/Full field) for 1, 5, and 10 stimuli at 100Hz shows a decreasing ration with stimuli LMM. Stimuli p = 1.4E-7. N=16 imaging fields from 11 slices from 3 mice. * = p<0.05, *** = p<0.001

### Functional consequences of rapid GABA diffusion

Modeling and confocal imaging results suggest that GABA diffuses from the site of release before GATs can remove it from the extracellular space. If true, we would predict that GABA released at one synapse could spill-over into neighboring synapses, and/or activate extra-synaptic (GABA_A_ or GABA_B_) receptors. To test the first possibility, we tested whether GABA released from one set of synapses could diffuse into neighboring synapses to induce hetero-synaptic GABA_A_ receptor desensitization^20^. First, we established a whole cell recording from a layer V pyramidal neuron to record IPSCs. Next, we electrically stimulated separate sets of inhibitory synapses using two bipolar concentric stimulators placed 200-350 µm medially and laterally from the recorded neuron^32^. The first stimulator delivered a single test pulse to evoke an IPSC. Then, we delivered 5 stimuli at 100 Hz to the second stimulator, to evoke a larger amount of GABA release, and then 100 ms later delivering another single test pulse to the first stimulator. We then isolated the IPSC evoked by the test pulse using a subtraction approach (Fig. 4A). If the preceding stimulus train caused heterosynaptic spill-over and desensitization of GABA_A_ receptors, this should reduce IPSC amplitude evoked by the test stimulation. Consistent with heterosynaptic desensitization, we found that the small burst of activity delivered to the second stimulator significantly reduced the amplitude of the IPSC evoked by the test pulse stimulation delivered to the first stimulator. This effect was enhanced by GATs inhibitors (NO-711 + SNAP5114; Fig. 4B/C) showing that GATs help constrain heterosynaptic signaling (LMM, Effect of pre-stimulus, Control IPSC: 0.87 ± 0.04, p = 0.038, SNAP5114/NO711: 0.73 ± 0.06, p=0.014, effect of SNAP5114/N0711 vs control, p= 0.032, N=12 cells from 5 mice). Lastly, to ensure that this paradigm did not cause a change in presynaptic GABA release, we repeated the experiment while quantifying GABA release with iGABASnFR1. Using this paradigm, evoked GABA release was unaffected by small bursts of activity at other inhibitory synapses (LMM ,ratio 1.00 ± 0.08, p = 0.48. N= 11 slices from 3 mice; Fig. 4D&E). This suggests that the GABA diffusion from one set of synapses can drive GABA_A_ receptor desensitization at neighboring synapses and demonstrates that even for brief stimulus trains, GABA can act heterosynaptically.

**Figure 4:**
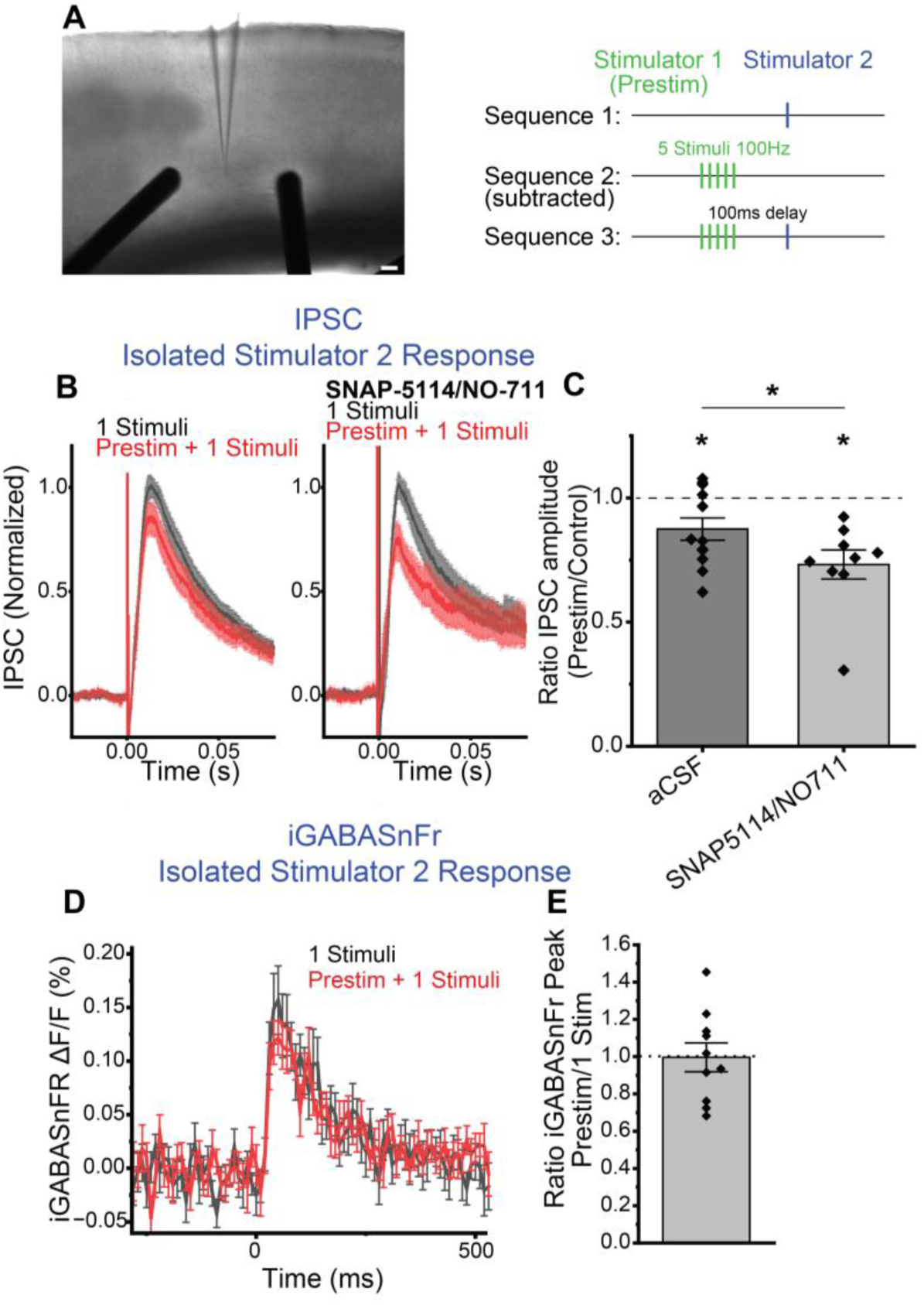
Local GABA diffusion reduces available receptors. **A)** Two stimulators are placed laterally in Layer V of the somatosensory cortex compared to the patch pyramidal neuron. On Stimulator 2, single stimuli are used to evoke IPSCs, with or without a preceding 5 stimuli 100Hz train on stimulator 1. IPSC responses to Stimulator 2 are isolated by subtracting traces. **B)** Isolated Stimulator 2 IPSCs show reduced amplitude when there is a preceding stimulus. Inhibiting GATs with SNAP5114 and NO711 enhances the effects of the prestimulus train quantified in **C)**. LMM, effect of prestimulus train Control p=0.0380, SNAP5114/NO711 p=0.014, effect of SNAP5114/NO711 p= 0.032 N=12 cells from 5 mice. Post-hoc test paired sample t-test p = 0.03, one sample t-test for aCSF p= 0.017, SNAP5114/NO711 p = 0.0019. **D)** iGABASnFR1 imaging with the same paradigm shows no effect of the presimulus train, suggesting no changes in presynaptic release. LMM p = 0.48. N= 11 slices from 3 mice. * = p<0.05

## Discussion

Here we show the first use of iGABASnFr to quantify GABA persistence in the extracellular space following synaptic release. Our results show that in the somatosensory cortex, the majority of synaptically released GABA rapidly diffuses away with only a small fraction removed by GATs. In the larger extracellular space, the persistence time of GABA is related to its local concentration. This is evinced by the long GABASnFR signal following stimulation (Fig. 1) and limited effect of GAT inhibitors both in epifluorescent imaging (Fig. 1), confocal imaging of GABA release sites (Fig. 3), and computational modeling (Fig. 2). Our findings build on previous modeling^23^, GABA_B_ measurements^26^, and more^10^ that suggest that GABA clearance by GATs is slow and that GABA can significantly diffuse in the extracellular space. There are small effects of GAT inhibition on synaptic iGABASnFr signals (Fig. 2, 3, S4), consistent with known GATs modulation of evoked IPSCs^5,15,22,23^. The effects of GAT inhibition are largely limited to evoked IPSCs and with no effect of GATs on spontaneous or unitary IPSCs^30^. We propose that during sparse synaptic release of GABA, as would occur during miniature or unitary IPSCs, diffusion is sufficient to restore extracellular GABA levels without engaging GATs. During a stimulus-evoked IPSC, however, there is sufficient local accumulation of GABA to engage GATs, but GAT transport is insufficient to constrain GABA diffusion. This suggests that the spatial organization of inhibitory synapses is crucial to the extracellular GABA dynamics, consistent with dense peri-somatic synapses being more sensitive to GAT inhibition than sparse distal synapses^8^. This spatiotemporal GABA summation in the extracellular space and is consistent with our finding that higher local GABA levels are associated with slowed clearance (Fig. 1).

Our findings also have important implications for the relationship between phasic and tonic inhibition. Assays of tonic GABA inhibition rely on the application of GABA_A_ receptor inhibitors, limiting our ability to monitor real-time changes. Our results suggest that, even for a single stimulus, GABA diffuses to and acts at neighboring synapses, rapidly transitioning from local phasic inhibition to broad tonic inhibition. This is partially masked in IPSC recordings by fast GABA_A_ receptor kinetics and desensitization. Similarly, our iGABASnFr data show that extracellular GABA accumulates extracellularly with increased stimuli number, consistent with continued GABAergic vesicle release^58,59^, but IPSC recordings (Fig. 1L/M) do not show the same magnitude of summation. This divergence between extracellular GABA and IPSC signaling can be explained by GABA_A_ receptor desensitization^21^. Together, this suggests that extracellular GABA dynamics rapidly blur the distinction between phasic and tonic GABA and represent a continuum based on the local spatiotemporal pattern of activity.

In summary, we use iGABASnFR imaging to identify novel and unexpected aspects of extracellular GABA dynamics that have important implications on synaptic integration, information processing, and the metabolic cost of sustaining GABA release during heightened activity. Our findings also suggest that synaptically released GABA rapidly diffuses to drive tonic GABAergic inhibition that is much more dynamic and spatially heterogenous than previously appreciated. New tools, including the inhibitory driving force sensor ORCHID^60^, genetically encoded voltage indicators, and neurotransmitter sensors could provide a completely new view of the spatiotemporal nature GABAergic inhibition and the role of GATs to provide largely slow, spatially broad GABA uptake. Our findings raise important questions about how GABAergic inhibition, which relies on fast-spiking neurons to drive temporal precision, can operate in a context of slow, spatially broad extracellular neurotransmission, suggesting we have much to learn about the fundamentals of inhibitory neurotransmission.

## Acknowledgments

This work was funded by the grants from the National Institute of Neurological Disorders and Stroke (R01NS127819 to M.A. and R01NS113499 to G.G.D.) and the National Institute of Aging (R01AG085666 to C.G.D), and by the Intramural Research Program of the National Institutes of Health (NIH; ZIANS003039 to J.S.D.). The contributions of the NIH author are considered Works of the United States Government. The findings and conclusions presented in this paper are those of the author and do not necessarily reflect the views of the NIH or the U.S. Department of Health and Human Services. We would like to thank Dr. Loren Looger and the GENIE Project for making plasmids and viruses available that were used in this study.

## Author contributions

Conceptualization, R. G., J.D., C.D., M.A.; Investigation, R.G., J.B., M.A; methodology, M.A., C.D.; formal analysis, R.G., M.A.; writing—original draft preparation, R.G., M.A., writing—review and editing, R.G., C.D., funding acquisition, R.G., C.D., M.A.; supervision, C.D., M.A.

---

**Figure Supplement 1:**
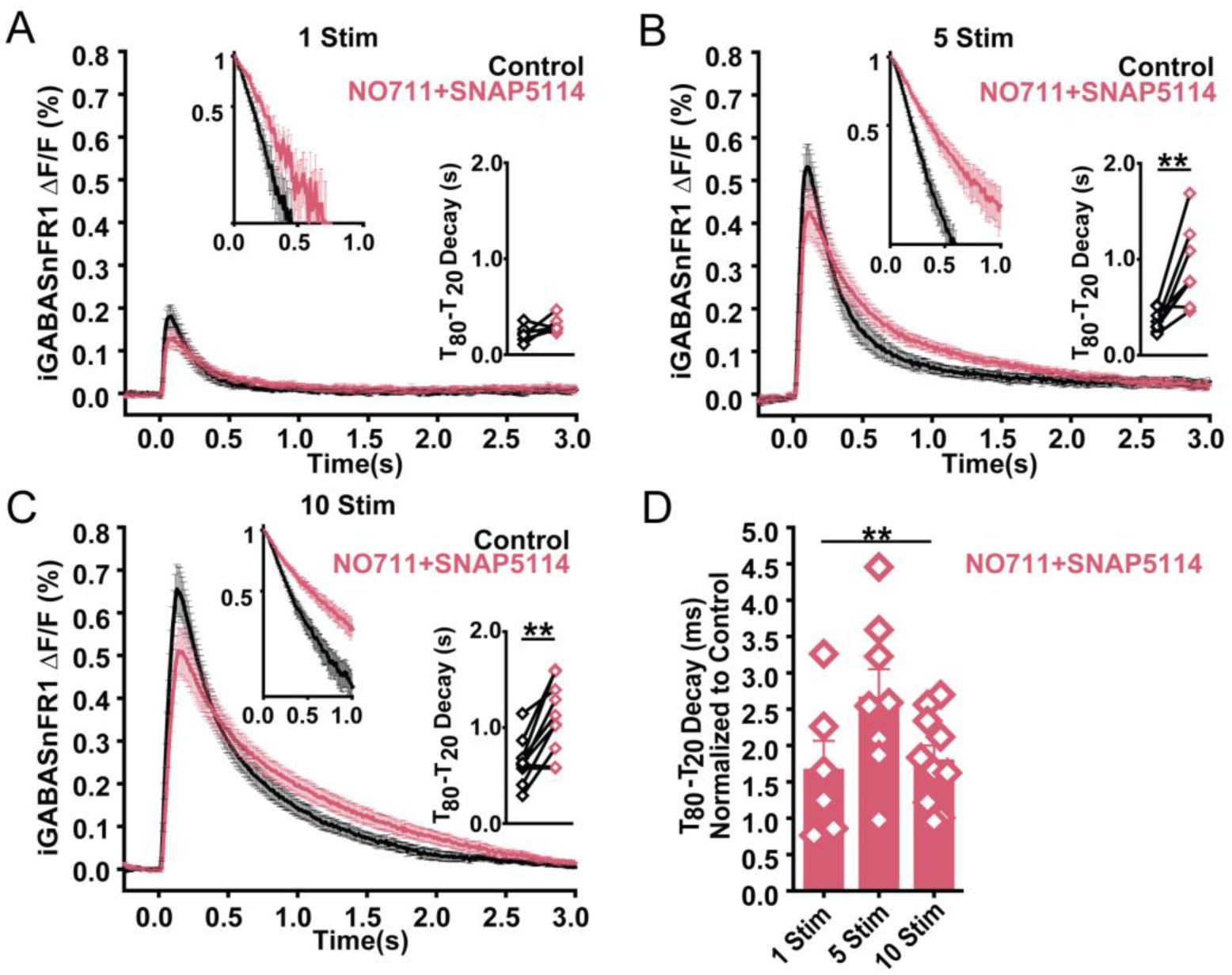
Inhibiting GATs slows iGABASnFr decays for all stimuli. **A)** Traces of iGABASnFR ΔF/F (%) over time (s) in layer V in response to 1 stimuli delivered at 100Hz in in both control and in GAT1+3 inhibitor group (treated with NO711 (25μM) and SNAP5114 (50μM)) with a semi log inset of the normalized fluorescent decay trace a line series inset of the T_80_-T_20_ Decay (s). Paired t-test (T_80_-T_20_ Decay (s)) control vs NO711+SNAP5114, p=0.15371. **B)** Traces of iGABASnFR ΔF/F (%) over time (s) in layer V in response to 5 stimuli delivered at 100Hz in in both control and in GAT1+3 inhibitor group (treated with NO711 (25μM) and SNAP5114 (50μM)) with a semi log inset of the normalized fluorescent decay trace a line series inset of the T_80_-T_20_ Decay (s). Paired t-test (T_80_-T_20_ Decay (s)) control vs NO711+SNAP5114, p=0.00497. **C)** Traces of iGABASnFR ΔF/F (%) over time (s) in layer V in response to 10 stimuli delivered at 100Hz in in both control and in GAT1+3 inhibitor group (treated with NO711 (25μM) and SNAP5114 (50μM)) with a semi log inset of the normalized fluorescent decay trace a line series inset of the T_80_-T_20_ Decay (s). Paired t-test (T_80_-T_20_ Decay (s)) control vs NO711+SNAP5114, p=0.00123. **D)** T_80_-T_20_ decay (ms) normalized to control. LMM **p<=0.00364. One Way ANOVA Tukey Test (T_80_-T_20_ Decay (ms) Normalized to Control), 1 Stim vs 5 Stim p=0.07567, 1Stim vs 10Stim **p=0.00681, and 5Stim vs 10Stim p=0.56956. N= 6 mice; 9 slices. ** = p<0.01

**Figure Supplement 2:**
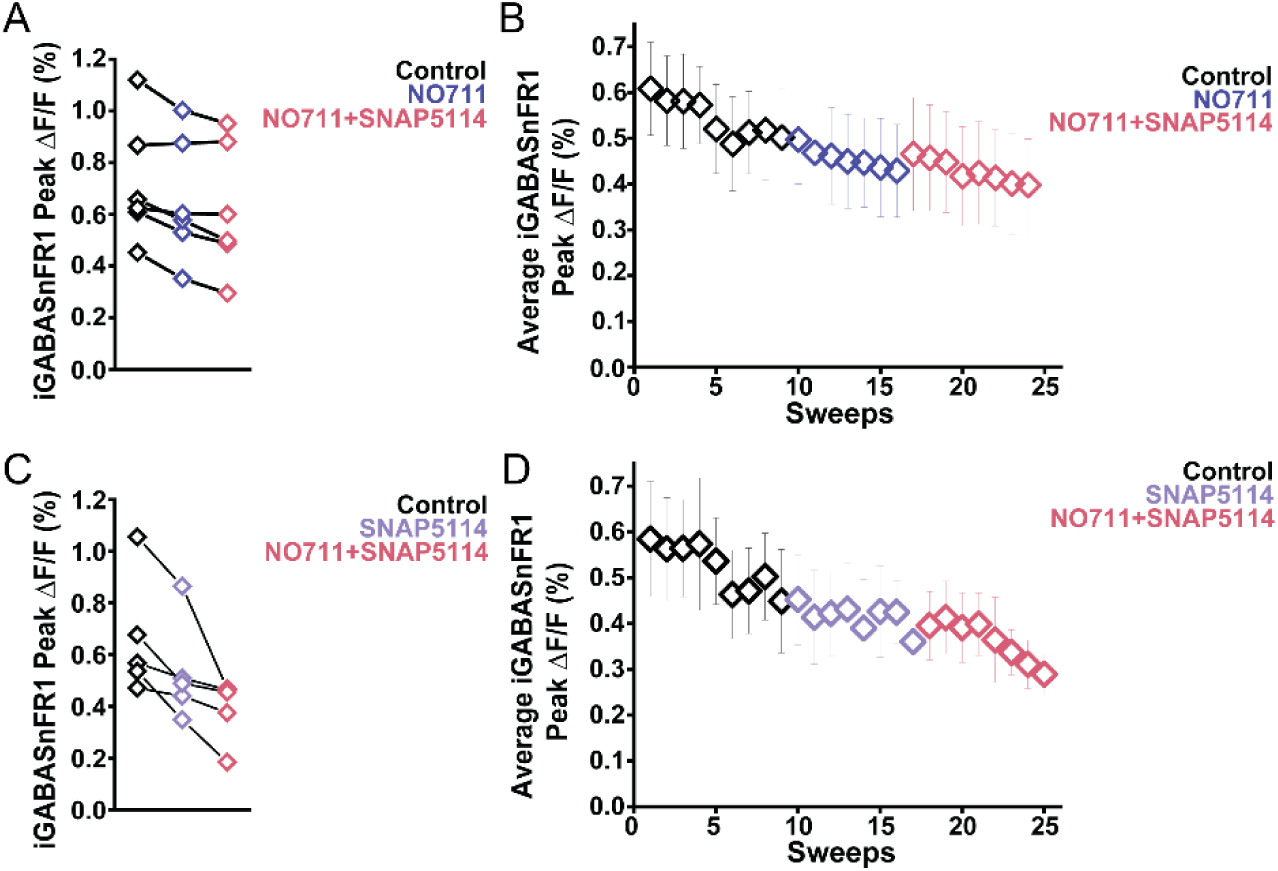
iGABASnFR1 bleaching during repeated imaging. Average iGABASnFR1 Peak ΔF/F (%) decreases regardless of drug condition as number of sweeps increase due to bleaching. **A)** iGABASnFR1 Peak ΔF/F (%) in response to 10 stimuli at 100Hz in control (treated with aCSF) and in NO711 (GAT1 inhibitor (25μM) and NO711+SNAP5114 (treated with GAT1 inhibitor (25μM) and GAT3 inhibitor (50μM)). NO-711 N= 2 mice; 5 slices. **B)** Average iGABASnFR1 Peak ΔF/F (%) vs number of sweeps in control, NO711, and NO711+SNAP5114 experimental sequence. LMM NS. NO-711 N= 2 mice; 5 slices. **C)** iGABASnFR1 Peak ΔF/F (%) in response to 10 stimuli at 100Hz in control (treated with aCSF) and in SNAP5114 (GAT3 inhibitor (50μM) and NO711+SNAP5114 (treated with GAT1 inhibitor (25μM) and GAT3 inhibitor (50μM)). SNAP5114 N= 5 mice; 5 slices. **D)** Average iGABASnFR1 Peak ΔF/F (%) vs number of sweeps in control, SNAP5114, and NO711+SNAP5114 experimental sequence. N= 5 mice; 5 slices.

**Figure Supplement 3:**
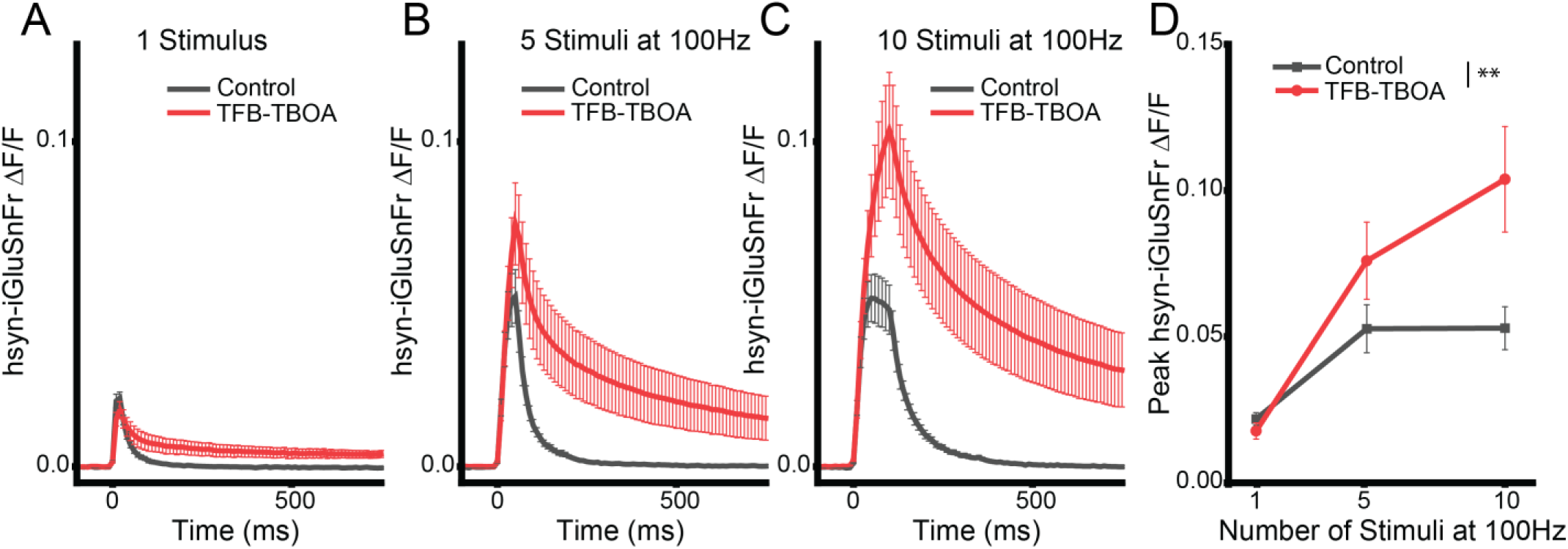
iGluSnFr shows rapid clearance during the stimulus train. Average hsyn-iGluSnFr traces in response to 1 **A)**, 5 stimuli **B)**, and 10 stimuli **C)** at 100Hz for control and with glutamate transporters inhibited with the pan EAAT inhibitor 1µM TFB-TBOA. The increase in peak fluorescence during the stimulus train represents active glutamate clearance during the stimulus train. **D)** Peak iGluSnFr fluorescence for control and TFB-TBOA traces shows increasing divergence, showing active transport. LMM Stimulus x TBOA interaction p = 0.0038. N= 10 slices from 3 mice.

**Supplementary Figure 4:**
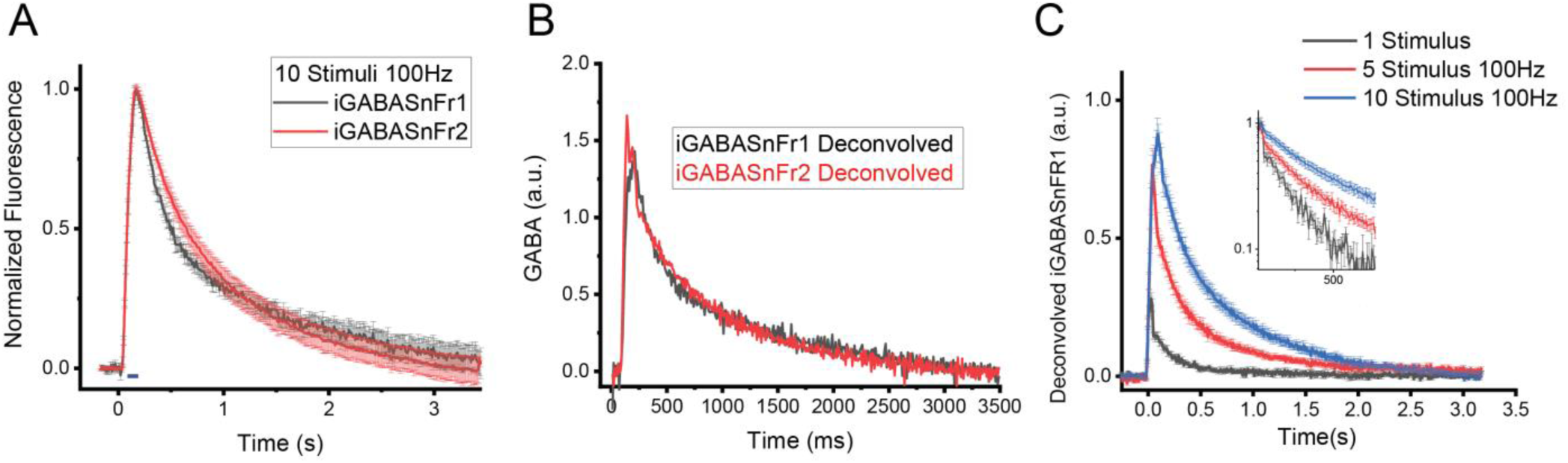
Deconvolving iGABASnFr fluorescence traces. **A)** Normalized, averaged iGABASnFr1 and iGABASnFr2 traces in response to 10 stimuli at 100Hz. Using the relative affinities of the two sensors along with the normalized traces, we derived the **C)** Average deconvolved traces for 10 stimuli at 100Hz for iGABASnFr1 and iGABASnFr2 shows similar kinetics for both sensors. **D)** Using the deconvolution function established in B, we deconvolved the average 1, 5, and 10 stimuli 100Hz iGABASnFR1 traces from Figure 1B.

**Supplementary Figure 5.**
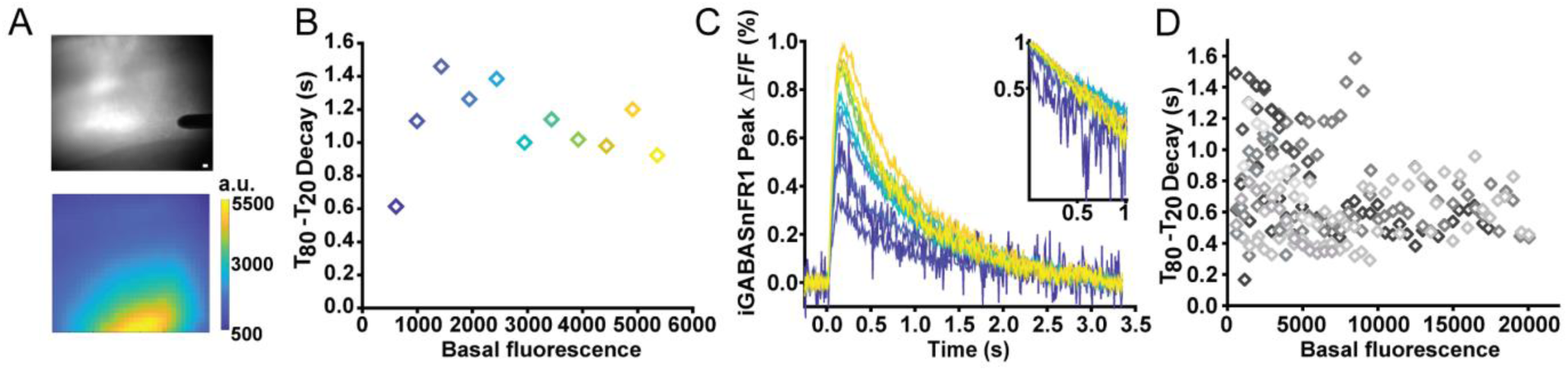
iGABASnFr decay uncorrelated with basal fluorescence intensity. **A)** Spatially averaged fluorescence response to 10 stimuli at 100Hz. Scale bar, 2.6μm. **B)** T_80_-T_20_ Decay (ms) vs basal fluorescence in response to 10 stimuli at 100Hz for 11 standard deviations for 1 slice with traces in **C.** in matching colors. **D)** T_80_-T_20_ Decay (ms) vs basal fluorescence in response to 10 stimuli at 100Hz for 10 slices. N=6 mice; 10 slices.

**Supplementary Figure 6.**
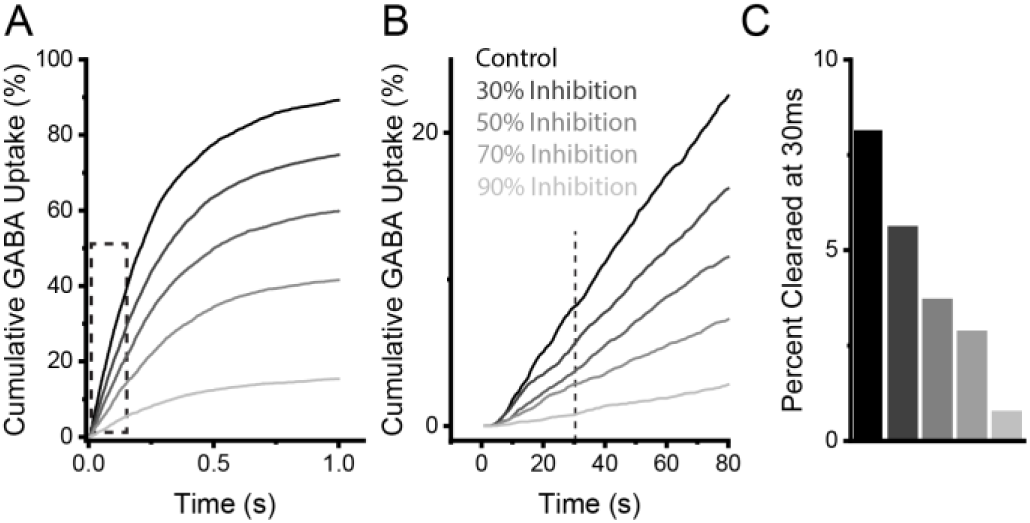
Simulation of cumulative GABA clearance by GATs. **A)** Modeling of cumulative GABA clearance with differing percent inhibition of GATs (30-90%) shows slowed GABA uptake. **B)** Magnified time-scale shows that GABA uptake starts within 10ms from release. **C)** Within the time-course of an IPSC (<30ms), a small fraction of GABA is cleared.

